# Arachidonic acid promotes the binding of 5-lipoxygenase on nanodiscs containing 5-lipoxygenase activating protein in the absence of calcium-ions

**DOI:** 10.1101/2020.01.22.915090

**Authors:** Ramakrishnan B. Kumar, Pasi Purhonen, Hans Hebert, Caroline Jegerschöld

## Abstract

Among the first steps in inflammation is the conversion of arachidonic acid (AA) stored in the cell membranes into leukotrienes. This occurs mainly in leukocytes and depends on the interaction of two proteins: 5-lipoxygenase (5LO), stored away from the nuclear membranes until use and 5-lipoxygenase activating protein (FLAP), a transmembrane, homotrimeric protein, constitutively present in nuclear membrane. We could earlier visualize the binding of 5LO to nanodiscs in the presence of Ca^2+^-ions by the use of transmission electron microscopy (TEM) on samples negatively stained by sodium phosphotungstate. In the absence of Ca^2+^-ions 5LO did not bind to the membrane. In the present communication, FLAP reconstituted in the nanodiscs which could be purified if the His-tag was located on the FLAP C-terminus but not the N-terminus. Our aim was to find out if 1) 5LO would bind in a Ca^2+^-dependent manner also when FLAP is present? 2) Would the substrate (AA) have effects on 5LO binding to FLAP-nanodiscs? TEM was used to assess the complex formation between 5LO and FLAP-nanodiscs along with, sucrose gradient purification, gel-electrophoresis and mass spectroscopy. It was found that presence of AA by itself induces complex formation in the absence of added calcium. This finding corroborates that AA is necessary for the complex formation and that a Ca^2+^-flush is mainly needed for the recruitment of 5LO to the membrane. Our results also showed that the addition of Ca^2+^-ions promoted binding of 5LO on the FLAP-nanodiscs as was also the case for nanodiscs without FLAP incorporated. In the absence of added substances no 5LO-FLAP complex was formed. Another finding is that the formation of a 5LO-FLAP complex appears to induce fragmentation of 5LO *in vitro*.

## Introduction

Leukotrienes (LTs) are formed initially through an interaction between 5LO (5-lipoxygenase) and FLAP (5-lipoxygenase activating protein), mainly in leukocytes. The LTs have essential roles in host defence as immunological actors in anti-inflammatory activities. Such activities may change into chronic inflammation states like asthma, allergy, rhinitis, rheumatoid arthritis and even lead to cancer and calls for extensive regulation mechanisms of the enzymes involved in LT formation (1–4). FLAP acts only as a scaffolding protein first presenting endogenous arachidonic acid (AA) to 5LO followed by stabilisation of the two-step oxidoreduction of AA to LTA4 (2).

As the FLAP/5LO proteins are almost exclusively expressed in myeloid cells (with life times as short as a few hours to days) which have to respond rapidly to inflammatory triggers, the two proteins are constitutively present and stored separately until needed, whereas AA is esterified and stored in the ER and nuclear membranes (5). FLAP is stored mainly in the nuclear membranes whereas 5LO is stored in the cytosol around actin or in chromatin in the nucleus (6, 7). It was found early on that the presence of an intact nuclear membrane may be essential for the combined activity of FLAP and 5LO as the increased product formation is observed only in intact cells (8). However, the recruitment of 5LO to the membrane does not require FLAP to be present, only Ca^2+^-ions are necessary for the translocation as shown in cells *e.g* (9, 10) as well as *in vitro e.g* (11, 12). It was proposed that 5LO binding is most productive in an initial stage and within 40 nm distance to FLAP according to a proximity ligand assay in cells (13, 14). At a later stage the LT production may be slowed, possibly due to a change in protein conformations or their tighter binding to each other (13, 14). The formation of a productive LT producing protein complex was shown to be dependent on the presence of AA, endogenously provided by the release from the membrane storage due to the action of cPLA2, cytosolic phospholipase A2 (10). A mutational analysis of 5LO showed that a triple lysine motif functions as a conformational switch regulator keeping the 5LO protected from turnover-based inactivation until a correct 5LO-FLAP interaction relieves the protection by the so called FY-cork for efficient LT synthesis (15). This interaction between 5LO and FLAP was shown to depend on Cys159 in that a mutation of this cysteine to serine inhibited the 5LO-FLAP complex formation (16). A cluster analysis (17) showed the complex formation could be relatively transient.

Here, the interaction of 5LO with FLAP was investigated using nanodiscs (ND) to assemble the complex. Nanodiscs are formed by phospholipids and two copies of a so called membrane scaffolding protein (MSP) (18, 19). This structure provides a well-defined template for studies of membrane bound – or membrane associated proteins as well as larger assemblies and is well suited for structure studies by cryoEM (20, 21).

The two questions we looked into were 1) would 5LO bind in a Ca^2+^-dependent manner also when FLAP is present? 2) Would the substrate (AA) have effects on 5LO binding to FLAP-nanodiscs? Is there a physical interaction in one or both of these cases? The latter may be hard to answer unless high resolution cryoEM could be applied. For the present communication negative stain electron microscopy was used together with gel electrophoresis and mass spectrometry to assess complex formation between 5LO and FLAP incorporated into nanodiscs. The results showed that presence of AA by itself induces complex formation in the absence of added calcium. Albeit *in vitro*, this finding corroborates that AA is necessary for the complex formation as proposed (10) and that a Ca^2+^-flush is mainly needed for the recruitment of 5LO and cPLA2 to the membrane. Our results also showed that the addition of Ca^2+^-ions promoted binding of 5LO on the FLAP-nanodiscs as was also the case for nanodiscs without FLAP incorporated (11). In the absence of added substances no 5LO-FLAP complex was formed. Another finding is that the formation of a 5LO-FLAP complex appears to induce fragmentation of 5LO.

## Materials and Methods

### Chemicals

If not otherwise stated, chemicals are from Sigma-Aldrich Co.

### Expression and purification of proteins

Human 5LO (UniProtKB: P09917) and FLAP (UniProtKB: P20292) was expressed in *E. coli* BL21-(DE3) (NEB). Expression of 5LO was performed in modified minimal medium (22) and the intrinsic ATP biding property of 5LO exploited during purification with ATP Agarose column (Sigma-Aldrich Co.) followed by gel filtration (23). The buffer was changed to 20 mM Tris pH 7.5, 100 mM NaCl, 2 mM EDTA, 1 mM FeSO4, 2 mM TCEP, 20 μg/ml catalase (11) for storage. 5LO was changed to the buffer 20 mM Tris pH 7.4, 100mM NaCl, and 0.5 mM EDTA to perform analysis and induce complex with 5LO. The concentration of 5LO in eluates was determined by Bradford assay (24).

The FLAP was expressed in modified LB medium and purified with engineered His-tag at the C-terminal (25). The FLAP was eluted in 20 mM Tris–HCl pH 7.4, 150 mM NaCl, 20 mM imidazole, 1 mM TCEP, 10% (v/v) glycerol and 0.1% (w/v) DDM. The buffer was changed to 20 mM Tris pH 7.4, 100mM NaCl, and 0.05% DDM for storage and analysis. The concentration was measured by the absorbance at 280 nm (ε = 56730 cm^-1^ M^-1^).

The MSP1E3D1 plasmid (with a His-tag as well as a TEV protease cleavage site, from Addgene, MA, USA) (26) was expressed and purified in *E. coli* (BL21-DE3) at the Protein Science Facility at Karolinska Institutet, Stockholm. The concentration of purified MSP1E3D1 (His-tag cleaved off) was determined by the absorbance at 280 nm (ε = 26600 cm^-1^ M^-1^).

### Preparation of FLAP-ND (FND)

The stock solution of 50 mM POPC (1-palmitoyl-2-oleoyl-sn-3-glycero-phosphatidylcholine) was prepared by firstly drying an appropriate amount of chloroform dissolved POPC (Avanti polar lipids, USA) under nitrogen followed by overnight removal of residual chloroform in a vacuum desiccator. This lipid cake was then resuspended in MSP standard buffer (25 mM Tris-HCl pH 7.5, 100 mM NaCl, 0.5 mM EDTA) supplemented with sodium cholate (Anatrace, USA) to a final concentration of 100 mM, so that the final POPC: sodium cholate molar ratio was 1:2. C-terminal His6-tagged FLAP was added to the resuspended lipids in a molar ratio of 1:70 (FLAP:POPC) and MSP1E3D1 was added in a molar ratio of 2:1 (MSP:FLAP). This reconstitution mixture was incubated on ice for one hour. The final concentration of sodium cholate was adjusted to 20 mM. Reconstitution of FLAP into nanodiscs was initiated by adding the Biobeads (Biorad) 0.5 mg/ml and incubating in a rotary incubator for 16 hours at 4°C (18). This reconstitution mixture was then purified by Ni-sepharose beads to fish our FND using the His-tag in FLAP. This was followed by clarification of eluates by centrifugation at 13000 x g for 10 min at 4°C and sample purification by gel-filtration using a GE Superdex 10/300 column equilibrated with MSP standard buffer. The peak fractions were collected and concentrated using centrifugal filters.

### Preparation of 5LO-FLAP-Nanodisc complexes

Although the order of mixing the components varied as stated in the text, the final concentration of the complexes always contained 0.6 μM 5LO and 0.6 μM FND in standard MSP buffer. The final concentration of additives was 50 μM AA or 0.5 mM free Ca^2+^-ions (adjusted with the EDTA present in the standard buffer). The total time used for mixing and incubations before loading on native gels were 10 minutes irrespective of the order of addition.

### Sucrose gradients

Two stock solutions of different concentration 5% (top solution) and 20% (bottom solution) were prepared by mixing corresponding amount of sucrose and dissolving them in a buffer containing 25 mM Tris-HCl pH 7.5, 100 mM NaCl, 10 mM β-mercaptoethanol. The gradient was prepared by filling the centrifuge tubes with 2.5 ml of top solution first followed by slowly injecting the bottom solution without disturbing the boundary layer. This tube was placed in a gradient master (Biocomp Gradient Master Model 106) and the program (47 secs, 86.5 degree, 25 rpm) was ran. Finally, 50 μl of sample was added on the top of the density gradient solution. The whole tube was loaded on to Sw50.1 rotor and ran in an ultracentrifuge at 50000 RPM for 16 hrs at 4 °C. After the run, the tube was carefully removed from the rotor and placed in a fractionator and fractions were collected (100 μl/tube) from the bottom of the tube.

### Electrophoresis

The native PAGE gel electrophoresis runs were performed in 4–16% Bis-Tris gel (NativePAGE™ Novex, Invitrogen). The samples (10 ng) were mixed with 4X Native PAGE Sample Buffer (Invitrogen) and loaded onto Bis-Tris gel. Blue native page was performed by using light cathode buffer (NativePAGE™ Running Buffer with 0.001% Coomassie G-250) in the cathode tank and NativePAGE™ Running Buffer as anode buffer. Samples ran on gel until the Coomassie front reached the end of the gel at a constant 150 V.

Denaturing PAGEs gel electrophoresis runs were performed in 10-20% Tris-glycine. The samples were mixed with 5X SDS PAGE loading buffer. After heating the sample for 10 minutes at 80 °C, it was loaded onto the wells and the samples ran until the dye front reaches the bottom of the gel at 150V.

The protein bands were visualized either by treating the gel with SilverQuest™ Silver Staining Kit (Invitrogen) or by Coomassie blue stain.

### In-gel Protein Digestion and Mass Spectrometry

Protein bands were excised manually from Coomassie- or silver-stained gels and in-gel digested using a MassPREP robotic protein-handling system (Waters, Millford, MA, USA). Gel pieces were destained twice with 100 μl 50 mM ammonium bicarbonate (Ambic) containing 50% acetonitrile at 40 °C for 10 min. Proteins were then reduced with 10 mM DTT in 100 mM Ambic for 30 min at 40 °C and alkylated with 55 mM iodoacetamide in 100 mM Ambic for 20 min at 40 °C followed by digestion with 0.3 μg trypsin (sequence grade, Promega, Madison, WI) in 50 mM Ambic for 5 h at 40 °C. The tryptic peptides were extracted with 1% formic acid in 2% acetonitrile, followed by 50% acetonitrile twice. The liquid was evaporated to dryness and the peptides were separated on an EASY-spray column connected to an EASY-nLC 1000 system (Thermo Scientific). The peptides were eluted in a 60 min gradient (from 5-26% of buffer B (2% acetonitrile, 0.1% formic acid) in 55 min and up to 95% of buffer B in 5 min) at a flow rate of 300 nL/min and analysed on a Fusion Orbitrap mass spectrometer (Thermo Scientific). The spectra were analysed using the Mascot search engine v.2.5.1 (Matrix Science Ltd., UK).

### Electron microscopy and image processing

3 μl of the sample was placed on a glow-discharged Cu-grid, coated with a C-film. Sample was incubated on the grid for 1 minute, blotted off with a filter paper, washed in two drops of water and stained 30 s in either 0.75% uranyl formate (UF) or 2% sodium phosphotungstate (NaPT, Polysciences Inc. USA), pH 7.5.

The grids were checked in a Jeol JEM-2100f microscope operated at 200 kV and images taken using defocus range 1.0-1.5 μm. NaPT-stained samples using TVIPS TemCam-415 4k × 4k CCD-camera (Tietz Video and Image Processing Systems GmbH, Germany) using nominal magnification 50,000x, which gives a calibrated value of 2.1 Å/pixel on the specimen level for CCD-images. Samples for complexes of 5LOFNDAA was taken from sucrose gradient fractions 9 and 10 and of 5LOFNDCa from fraction 9. Images from UF-stained samples were collected on a DE-20 camera (Direct Electron, LP, USA) at a nominal magnification of 40,000x, resulting in a sampling distance of 1.54 Å/pixel. Each image on the DE-20 detector was exposed for 2 s using a frame rate of 20 frames per second. Before image processing, the collected frames were first motion corrected. The first two frames for each image were discarded due to drift.

Image processing was performed using programs from the EMAN2.12 package (27). The input particle sets for NaPT-stained samples were composed of 4541 (FND), 6305 (5LOFNDAA) and 5714 (5LOFNDCa) particles. For UF-stained samples 2D classifications particle sets were of sizes 3081 (FND), 3705 (5LOFNDAA) and 3201 (5LOFNDCa). Initial models were created from 2D-class averages of the NaPT stained particles. 3D-models were visualised using UCSF Chimera (28).

## Results

### Separation of nanodiscs containing FLAP from empty nanodiscs

In earlier experiments regarding Ca^2+^-dependent binding of 5LO on nanodiscs (11), native gel electrophoresis showed a band corresponding to 5LO in complex with so called “empty” nanodiscs (END), i.e. nanodiscs containing only phospholipids (*i.e*. POPC) and MSP. In the present communication, nanodiscs containing FLAP (FND) and also POPC were used in order to assess binding characteristics of 5LO to nanodiscs when FLAP is present in the membrane. These FNDs were prepared using C-terminally tagged FLAP (FLAP-C-His6). Nanodiscs containing FLAP could be purified by affinity chromatography using a C-terminal tag on FLAP but not when the tag was at the N-terminus (data not shown). This may be due to the FLAP N-terminus being localized deeper in the trimer or in the surrounding membrane as compared to the C-terminus. After affinity chromatography to separate the FLAP-containing nanodiscs from ENDs, size exclusion chromatography (SEC) was carried out. 10 ng of a representative SEC fraction loaded on a native gel (Fig. 1A, left lane) shows a band at somewhat higher Mw than 242 kDa which corresponds to where END is usually observed.

**Fig 1.**
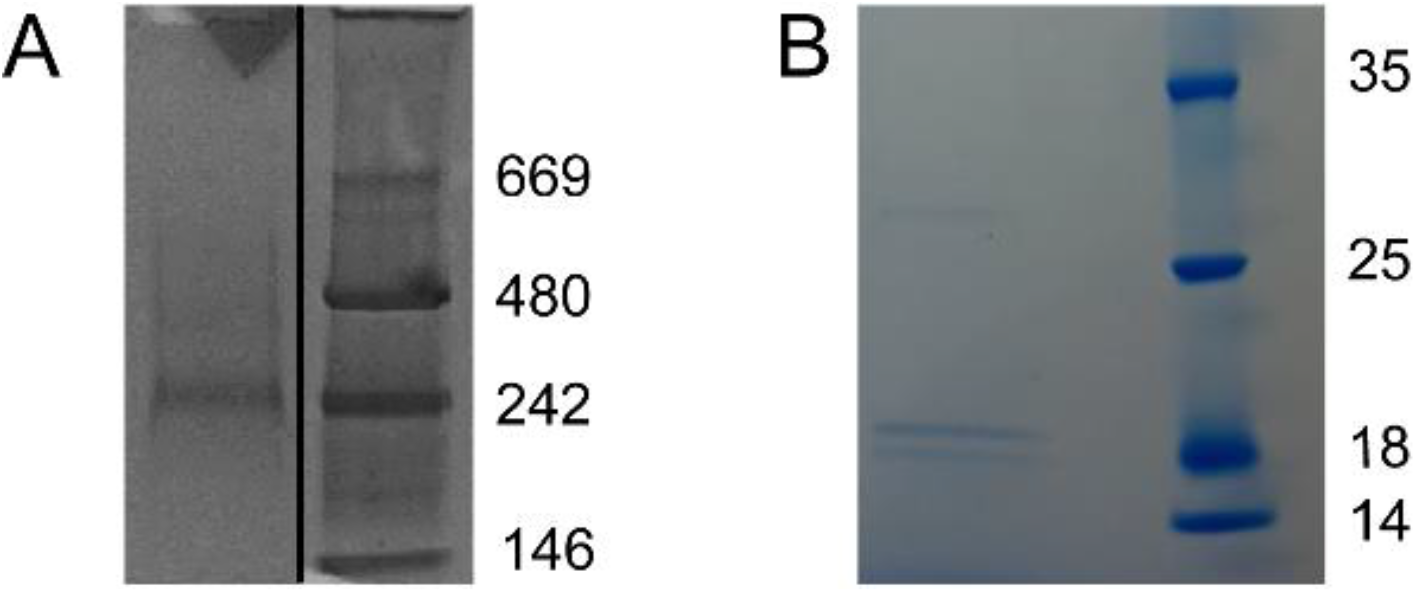
FLAP can be reconstituted into nanodiscs. (A) In the native gel, a band corresponding to purified FND (left lane) runs slightly slower than 242 kDa (see marker lane to the right) where END runs (11). (B) A sample of FND denatured by SDS was loaded in the left lane and shows bands corresponding to MSP1E3D1 (29.9 kDa) and FLAP-C-His6 (18.5 kDa). To make the figure, relevant lanes were cut in the original gel image and rearranged in (A) and a section was cut out from the original gel image in (B), see S1_Raw_images.pdf.

This is not surprising as FLAP is expected to protrude only slightly from the membrane (29, 30), see Fig. 2. Hence, the size and shape of the FND is almost identical to the END when they are both made with MSP1E3D1 (18). A part of the SEC fraction sample loaded in Fig. 1A was denatured by SDS and loaded on a denaturing gel as shown in Fig. 1B, left lane. Bands corresponding to the expected proteins can be observed; MSP1E3D1 at 29.9 kDa and FLAP-C-His6 at 18.5 kDa. The band at around 18 kDa is occasionally observed in samples where FLAP has been denatured by SDS.

**Fig 2.**
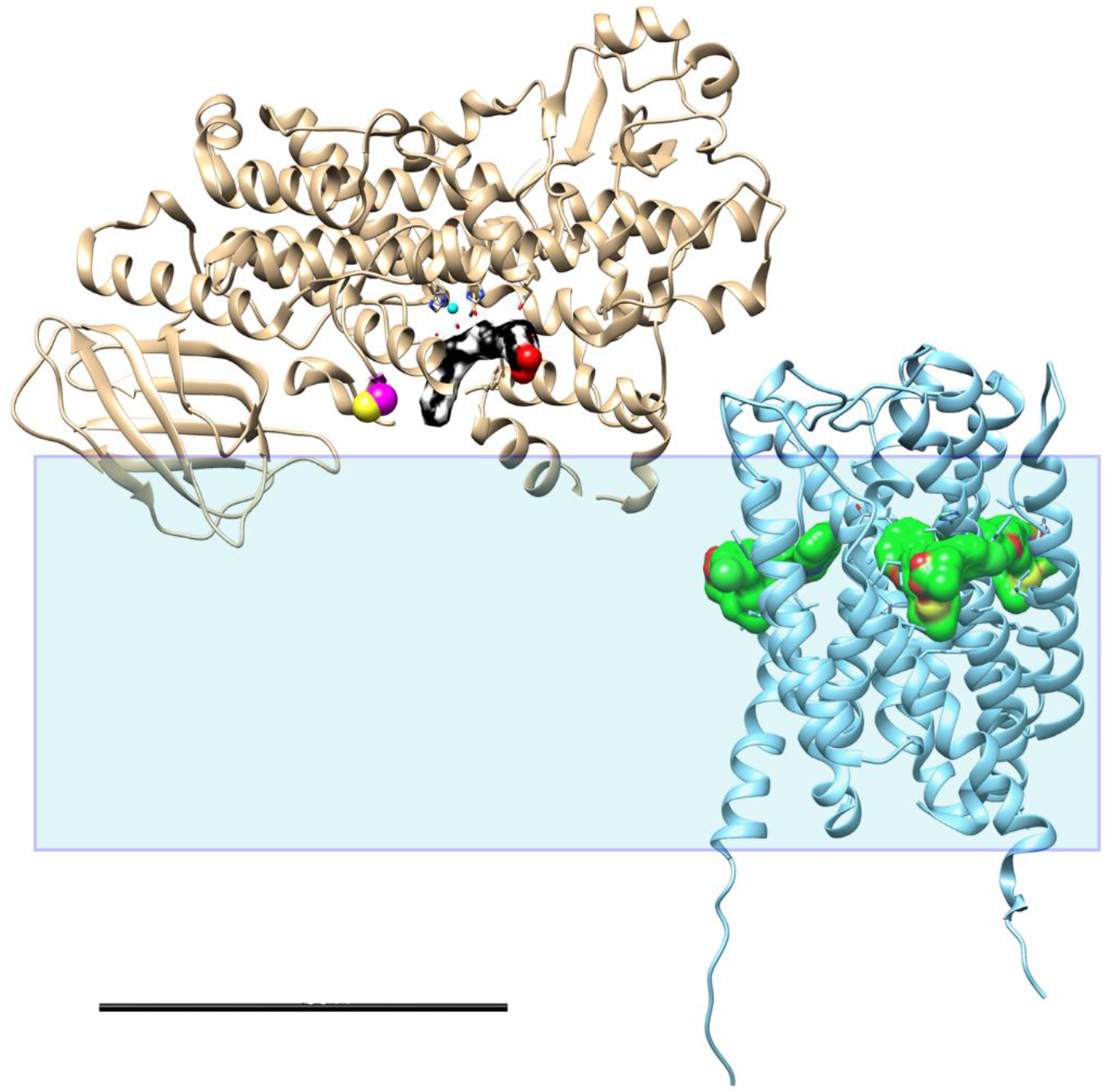
Schematic view of 5LO and FLAP embedded in the membrane. Ribbon models of 5LO (left, PDB ID: 3V99) and FLAP (right, PDB ID: 2Q7M) shown in relation to the membrane, highly schematically represented by the transparent rectangle which is 5 nm thick. NB! It is not known how deep 5LO penetrates the membrane or how much FLAP protrudes from the membrane. The substrate AA in 5LO (black carbon atoms, red oxygens) and three inhibitor molecules (MK-591) in FLAP (green carbon atoms, red oxygens, yellow sulphurs) are shown in surface representation. In 5LO the non-haem iron (cyan) is surrounded by three histidines and water whereas the Cys159 shown to mediate binding to FLAP (16) is shown as spheres (yellow, magenta). Scale bar 5 nm

### The presence of AA is sufficient to induce binding of 5LO on the FLAP-nanodiscs

In earlier work it was shown that 5LO could form a complex with nanodiscs, but only if calcium ions were added to the solution of 5LO and nanodiscs (11). This complex, 5LOEND, showed as a band on a native gel as well as in denaturing SDS-PAGE. Addition of AA to the complex lead to a strongly increased product formation whereas substrate addition to a solution of 5LO and ND in the absence of Ca^2+^-ions lead to the basal low rate of product formation similar to the rate catalysed by unbound 5LO. Hence, it appeared AA alone did not induce binding of 5LO on the END.

In the present work, we analysed the effect of AA and Ca^2+^ with nanodiscs containing FLAP and 5LO in denaturing gels as well as in native page gels. 5LO was present but also either calcium ions or the substrate, AA. Since FLAP is present in the nanodiscs and is discussed as an activator that presents AA to 5LO, it was interesting to investigate effects AA could have on its own, i.e. in the absence of calcium ions. Interestingly, we found that AA is sufficient to bind 5LO to the FLAP-nanodiscs. Several gel analyses from these different constellations of complexes, indicated that a 5LOFND complex could be formed either in the presence of calcium-ions (5LOFNDCa) or induced by AA (5LOFNDAA). However, observations of complex formation due to either Ca^2+^ or AA on the same native gel (for examples, see Figure 1 in the S2_file.pdf) were very hard to obtain and it does appear as if the protocol for sample preparation matters to some extent. Another method of observation was needed. It was also an aim of this project to obtain larger amounts of the complexes for future cryo-electron microscopy imaging and stabilised to allow for some storage before use. Although sucrose concentrations for cryo electron microscopy imaging need to be kept very low (31, 32), sucrose gradient separation of complexes followed by denaturing SDS page showed very promising results for the stability of the complexes.

### FLAP and 5LO in complexes purified by sucrose gradient separation after addition of either Ca^2+^ or AA

As indicated in the figure legend, seven fractions from each centrifuge tube were loaded on the denaturing SDS-PAGE gel shown in Fig. 3. For the Ca^2+^ -induced complex, *e. g*. fraction C9, clearly contained all three proteins expected in the 5LOFND complex, 5LO, FLAP and the MSP1E3D1. Also a breakdown product of 5LO appears to be present (ca 40 kDa).

**Fig 3.**
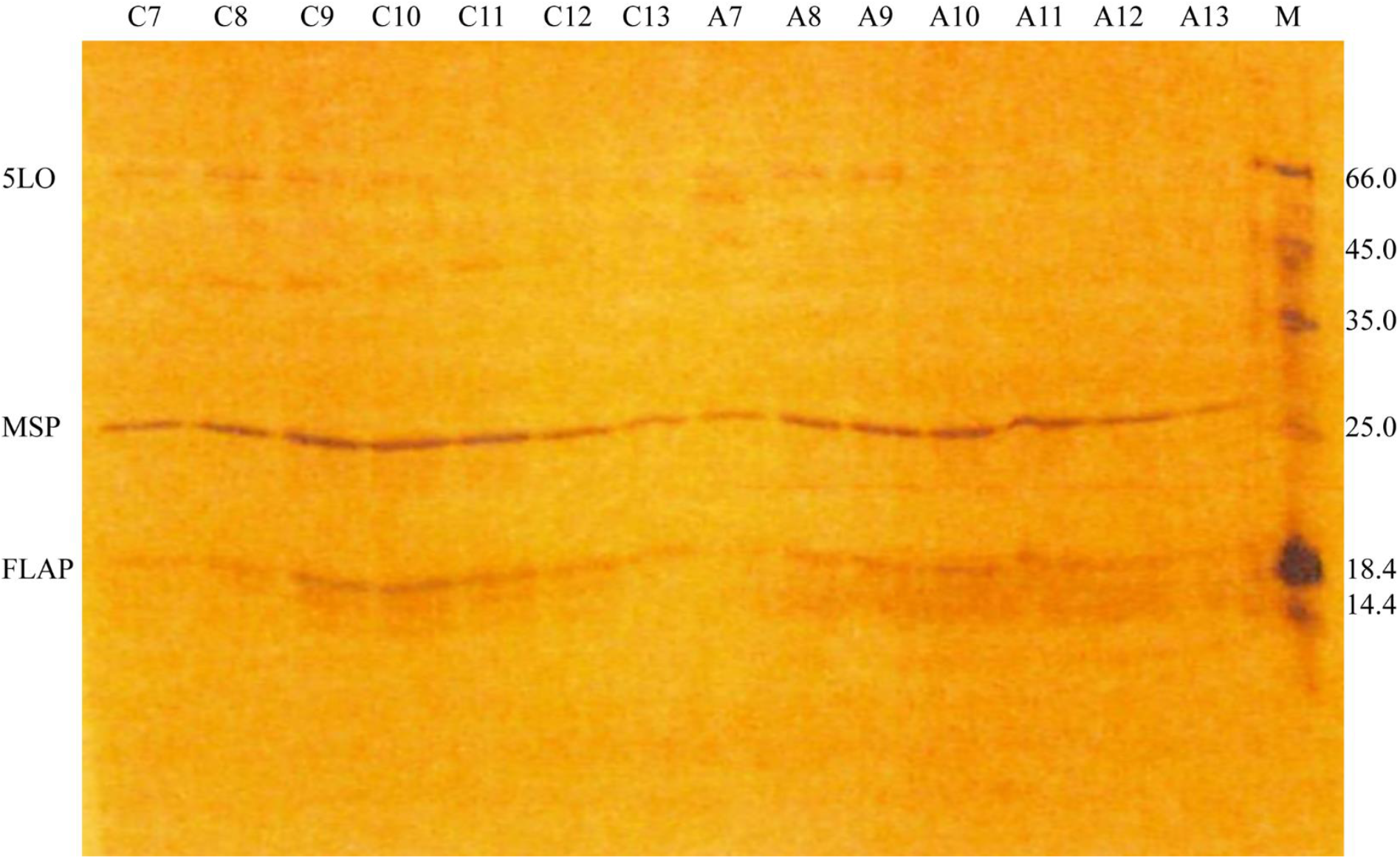
Either Ca^2+^ or AA can induce 5LOFND-complex formation as shown by SDS-PAGE denaturing gel electrophoresis after sucrose gradient separation. For the Ca^2+^ induced 5LOFNDCa-complex, seven fractions from the sucrose gradient were loaded on the gel (Lanes C7-C13). Out of these, fraction C9 shows the most intense bands of FLAP, MSP and 5LO. For the AA induced 5LOFNDAA-complex, seven fractions from the sucrose gradient were loaded on the gel (Lanes (A7-A13). Out of these, fraction A9 shows the most intense bands of FLAP, MSP and 5LO.

For the AA induced complex formation, the fractionation showed clear presence of the three proteins in fraction A9. Here, 5LO appears to remain intact.

In conclusion, the sucrose gradient fractionation shows that the complex of 5LO with nanodiscs containing FLAP is formed either in the presence of calcium ions or is induced by arachidonic acid.

### The complex induced by AA contained both 5LO and FLAP as detected by mass spectrometry

From a native gel, similar to those in Figure 1 in the S2_file.pdf, bands were cut and subjected to mass-spectrometry analysis. The result shows that the band from a complex induced by AA, contained the MSP, 5LO and FLAP (Fig. 4A). However, the band cut from the lane representing the Ca^2+^-induced complex on the same gel, did not contain any of the proteins in detectable amounts (Fig. 4B).

**Fig 4.**
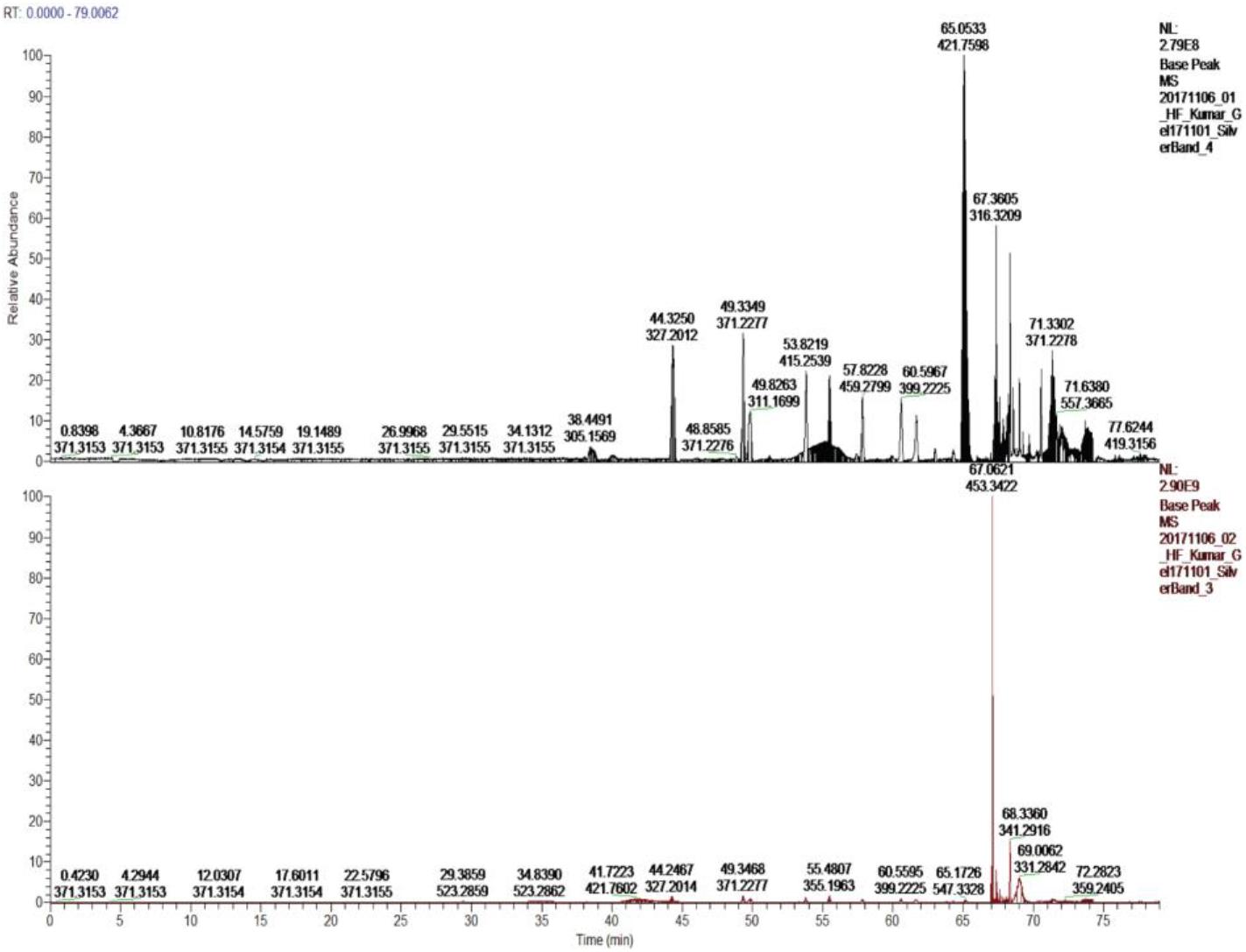
The arachidonic acid induced complex formation was detected with Mass-spectrometry. Samples for MS were obtained from a native gel similar to those presented in Figure 1 in the S2_file.pdf. Bands were cut from the gel and treated for mass-spectrometric analysis as described in the experimental section. In A) all three proteins, FLAP, 5LO and MSP1E3D1 are present whereas in B) none of the three proteins are present.

The latter did not make sense to us, at least the FND proteins should be identified by the mass-spectrometry. Bands were therefore cut from three other silver stained native gels and submitted for MS-analysis. Here, all bands were shown to contain both MSP1E3D1 and FLAP but not 5LO in any case. Inspection of Figure 1 in the S2_file.pdf shows that the complexes may show as smeared bands and one issue in cutting bands from native gels may be how much of the smeared bands to take and where to cut. The detection limit is another issue and for controls where 5LO or FND had been run on native gels and stained with silver, for example 5LO could not be detected by MS although the band was visible on the gel. In the control FND, both FLAP and the scaffolding protein could be detected (not shown).

### Negative stain electron microscopy of complexes formed by 5LO bound to FND

Samples were taken from incubations of FND and 5LO with Ca^2+^ or AA followed by fractionation on a sucrose gradient. A control was made from an END preparation as well as from FND without added 5LO. NaPT staining caused a heavy stacking of empty NDs as previously reported (11, 33). Stacking was, however, not observed when FND (Fig. 5A) or preparations of FND with Ca^2+^ or AA together with 5LO were stained with NaPT (Fig. 5B and C). The prevention of stacking indicates that the nanodisc lipid surface is not free to interact with another nanodisc surface. Instead, insertion of a membrane protein (FLAP) or binding of a membrane-bound protein (5LO) to the nanodiscs or both prevent the interaction between the nanodisc surfaces (33, 34).

**Fig 5.**
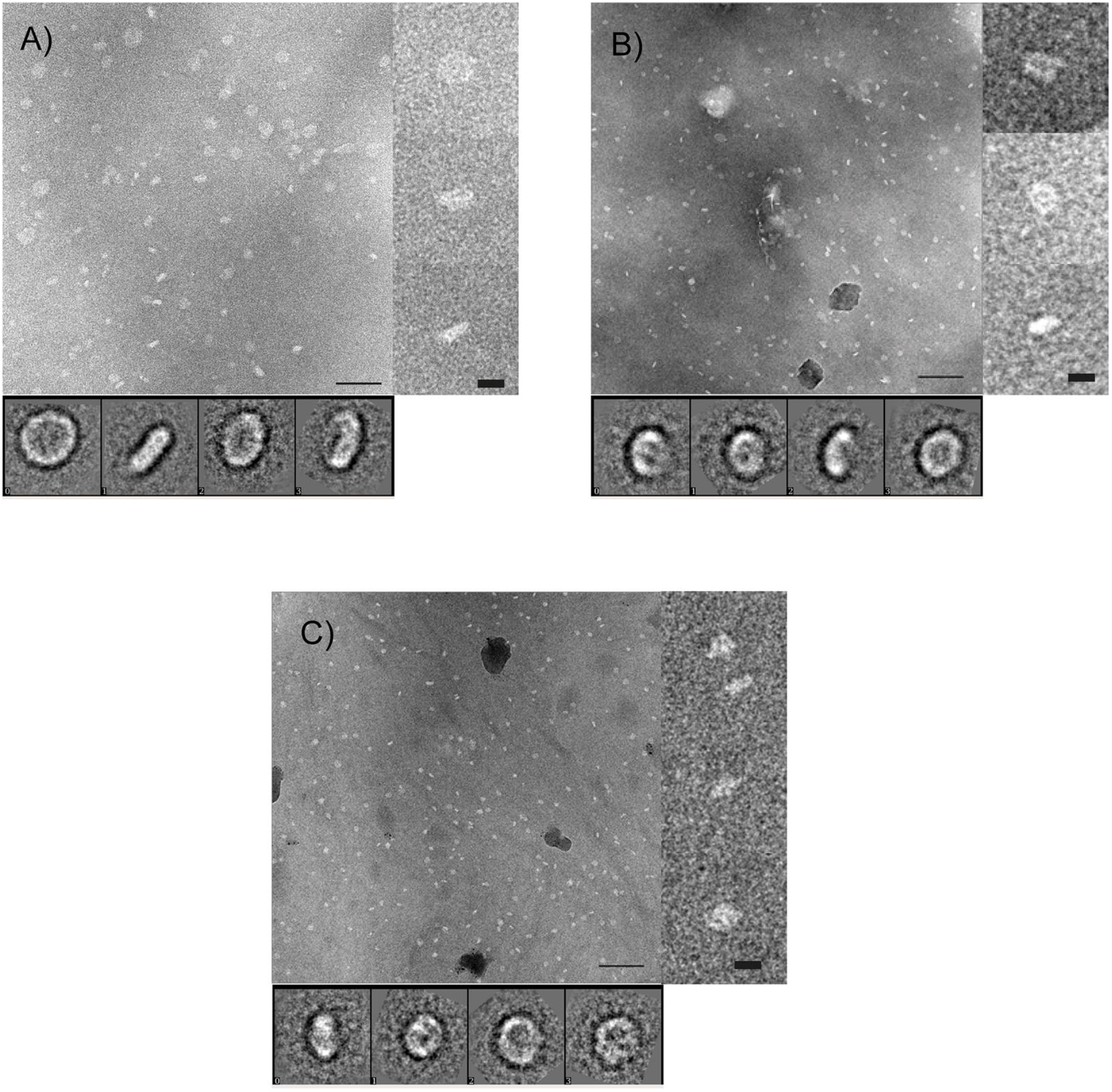
Negative stain TEM indicates formation of 5LO-FLAP nanodisc complexes. NaPT stained (2 %) images of FND:s in the presence or absence of 5LO. The image in A) shows FND:s in the absence of 5LO. These control FND samples were not purified on a sucrose gradient, hence larger particles are present compared to the images in B and C. Here the FND:s were mixed with 5LO and either B) arachidonic acid or C) calcium ions. The samples were collected after sucrose gradient separation. Insets at right show magnified typical views of the particles. Insets below show averages after 2D-classification (27). No stacking is observed in any image. Scale bar 100 nm, for insets to the right 10 nm and of the 2D classes the box side 26.9 nm.

As uranyl formate is the stain with the smallest grain size and thereby could provide a higher resolution, we collected images of UF-stained samples for further analysis (Figure 2 in the S2_file.pdf) (35). In 2D class averages from UF-stained FND samples the presence of FLAP might be seen by the presence of “holes” or features resembling stripes across the surface (Figure 2A in the S2_file.pdf). All views observed however, are perpendicular (“top views”) or slightly tilted relatively to the normal of the membrane plane. The FND preparation used had not been purified by sucrose gradient treatment hence showing some variation in disc size. The diameters of the top views in 2D-classes span between 10 nm and 20 nm. Thereby one or two FLAP proteins could be inserted per disc (Figure 2A in the S2_file.pdf).

Some differences in the 2-D classes are seen when 5LO was incubated with FND in the presence of AA or Ca^2+^ as compared to plain FND samples (Figure 2B and 2C respectively in the S2_file.pdf). Some classes of the UF stained complexes seem more tilted and blurry on the side as if there is an object *i.e*. the catalytic domain of 5LO on top of the disc. This is for example observed in class averages 18, 23 and 26 in Figure 2B in the S2_file.pdf showing AA induced complexes or class-averages number 18, 20 and 28 for calcium ion induced complexes (Figure 2C in the S2_file.pdf). These types of blur are not present in the control FND (Figure 2A in the S2_file.pdf) whereas some classes where we expect the complexes in Figure 2B and 2C in the S2_file.pdf instead resemble FND without a 5LO bound.

As use of UF mainly gave top views of the ND-samples on carbon film, while an increased number of side views were observed with NaPT-stain, we collected new data sets from NaPT-stained samples obtained from sucrose gradient fractions. All three sets (FND, 5LOFNDAA and 5LOFNDCa) show side views in their respective 2D class averages (Fig. 5A-C). Generally the results resemble those observed with UF; FND shows size variation of the discs (Figure 5A) and presence of 5LO can be suspected in some side/tilted views of the complex (Figure 5B and C).

Although NaPT generally produced lower contrast than uranyl stained samples, we collected data-sets as the presence of tilted views made it possible to calculate 3-D reconstructions. FND showed a disc-like 3D-volume that could allocate one or two FLAP proteins (Fig. 6 yellow). The complexes of 5LOFND in presence of AA or Ca^2+^ were similar, both showing an extra density on top of one side of the disc as compared to FND (Fig. 6). This should result from the membrane-bound 5LO. The extra density had also approximately the size of the catalytic domain of 5LO as was shown by docking the atomic structure of human 5LO (PDB ID 3O8Y) (36) manually into the extra density (Fig. 6C and D).

**Fig 6.**
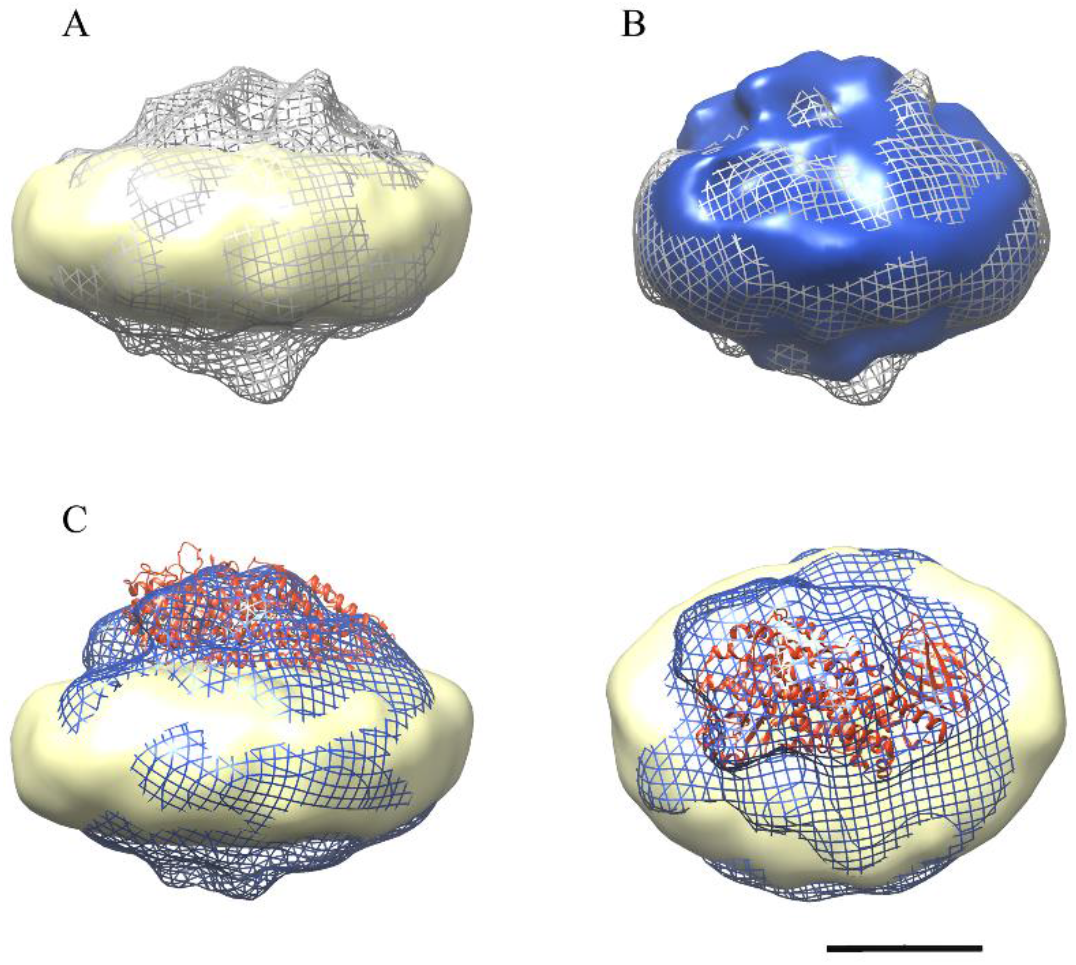
3D-reconstructions of NaPT-stained particle sets. A) Complex 5LOFNDAA (gray mesh) and FND (yellow). B) Complexes 5LOFNDCa (blue) and 5LOFNDAA (gray mesh) show similar densities. C) Side and top views of the complex 5LOFNDCa (blue mesh) and FND (yellow). The atomic structure of 5LO (PDB ID:3O8Y, red ribbon) was manually docked to the map. Scale bar 5 nm.

Although the size of the FND part of the 3D-reconstructions of complexes roughly corresponded to that of plain FND, they had a bit more compact appearance. This difference is probably induced by the binding of 5LO to lipid membrane, which slightly changes the form of the disc. Our results also suggest small structural differences between the two complexes, 5LOFNDAA and 5LOFNDCa. First, there are slight differences within the 5LO density on top of the disc, and secondly at the opposite side of the disc, where the volume in 5LOFNDAA is more bulging (Fig. 6B). It is here where FLAP can be expected to protrude from the membrane. However a higher resolution cryo-EM reconstruction is required to give definite answers of the structural changes between the complexes.

## Discussion

In a previous report we asked: would 5LO bind to the membrane in our particular system, the nanodisc? Would there be differences compared to the well-established liposomes for which only calcium ions are necessary for the translocation to and binding on the liposomes? These two platforms for *in vitro* studies of 5LO binding to phospholipid membranes proved to be very similar with the benefit that the nanodisc complexes can easily be visualized by transmission electron microscopy. For the present communication we investigated whether interactions between FLAP and 5LO would be possible to observe using nanodiscs containing FLAP. Would the presence of AA be essential for binding 5LO to the FND?

An interesting situation underlines these questions: It is known that FLAP is not necessary for 5LO to bind to the membrane *in vivo* in cells, on the other hand the sole known function of FLAP is to activate 5LO to yield high levels of LTA4 from endogenous AA stored in and released from the nuclear membranes or the ER. Our observations indicate a physical interaction and that this interaction could lead to damage and degradation of 5LO under certain conditions. This shows as split bands or even as fragments on gels.

### Either Ca^2+^ or AA can induce 5LOFND-complex formation. Stability aspects of the complexes

A prerequisite for the present communication was that FLAP could be incorporated in nanodiscs of the same size as used for the END:s in our earlier work, *i.e*. using MSP1E3D1 as the membrane scaffold protein which yields nanodiscs with an outer diameter of 12.8 nm (19, 37). Furthermore, negative staining with sodium phosphotungstate leads to a variety of orientations of the nanodiscs allowing for a 3D reconstruction to be made (Fig. 5 and 6). The ca 90 nm^2^ membrane area is just large enough (18, 37) to house both one FLAP and one 5LO which measures about 3.6 nm in diameter and 4.5 x 9 nm respectively (29, 36). Although a His-tag located on the N-terminus of FLAP leads to reasonable yields of detergent-purified protein after bacterial expression (not shown), only a His-tag located on the C-terminus on FLAP served its purpose in the purification of nanodiscs containing FLAP (Fig. 1). Examination of the FLAP structure shows that the N-termini turn towards the centre of the trimer and are located in the hydrophobic region of FLAP (Fig. 2, right). These do not protrude from the membrane in contrast to the C-terminal 20 residues in the human FLAP which are partly unstructured in the crystal (29) (Fig. 2, right). It would be interesting to see if these would become structured under other conditions e.g. close to other proteins (10, 17, 38) or would remain disordered. Simply the location of FLAP in a lipid environment could have induced order, however, specific protrusions from organised C-terminal FLAP extensions in the non-complexed FND:s could not be observed at our resolution in negative stain (Fig. 6, yellow).

Similar to the earlier results with 5LO and ENDs (11), calcium ions could promote the binding of 5LO on the FNDs as seen by negative stain electron microscopy and by electrophoresis. As observed in the micrographs (Fig. 5) and after data processing a 3D reconstruction of 5LOFNDCa shows densities (Fig. 6, blue) not present in the FLAP-nanodiscs (Fig. 6, yellow). On several native gels however, we failed to observe the 5LOFNDCa-complex. This may of course be due to a lower yield of complex-formation before the electrophoresis. A reason for this could be the smaller available phospholipid area and/or a possible steric hindrance due to FLAP. Yet, when the 5LOFNDCa samples were purified on a stabilising sucrose gradient followed by analysis of the fraction representing the complex on a denaturing gel, bands representing FLAP, MSP1E3D1 and 5LO were present indicating complexes containing all three proteins (Fig. 3). These complexes can be observed directly in the electron microscopy images in Fig. 5B and C. Therefore, likely due to the lack of calcium ions to stabilise the complex, the complexes fell apart during the gel electrophoresis (Figure 1 in the S2_file.pdf). Another observation can be made regarding the possible fragmentation of 5LO (Fig. 3) under some conditions as will be discussed below.

The presence of AA induced complexes of 5LOFNDAA despite the absence of calcium-ions as shown by sucrose gradient separation and electron microscopy (Fig 3 and 6). Furthermore, mass spectrometry also identified all three proteins (Fig. 4A). The AA-induced complex did show instability and split 5LO-bands similar to the Ca^2+^-complex as discussed above for the native gels (Figure 1 in the S2_file.pdf). However, for complexes separated by a sucrose gradient it appears as if 5LO remains intact under these conditions.

As discussed in (39) calcium dependent LT formation may be circumvented under cellular stress conditions implying that 5LO translocation has occurred independent of calcium flushes. In fact, it was shown that the C-terminal catalytic domain of 5LO shows membrane binding capability in the absence of the N-terminal domain which is responsible for the Ca^2+^ - dependent embedding in the nuclear membranes (40). Furthermore, the concentration of AA in different cells can be rather high even in un-activated states and partitions strongly to the membranes (5) and as discussed in (41, 42) AA may be provided via transcellular routes. Addition of AA after inhibition of cPLA2 activity rescued the formation of 5LOFLAP complex formation (13). In the same publication it was proposed that FLAP is a regulator which first interacts in a loser, flexible fashion with 5LO followed by tightening the interaction to a complex aimed for termination of the activity. Cys159 could be involved in this although the cytosol is a reducing environment (16). In polymorphonuclear leukocytes a conformational change in the N-and C-termini of FLAP was observed on binding of AA using Fluorescence Lifetime Imaging Microscopy (10). In the crystal structure of FLAP three inhibitor molecules occupy the three active sites (Fig. 2, right, green). The inhibitor binding sites were proposed to overlap with the AA-binding sites (43, 44). It is not known whether FLAP has a 3/3 sites reactivity like LTC4 Synthase (45) or 1/3 sites reactivity like for MGST1 and MGST2 (46, 47). 5LO has one active site per unit located in its C-terminal domain (Fig. 2 left) (36). It would be interesting to use a larger nanodisc or vesicles to investigate whether three 5LO would bind to one FLAP. On the other hand, the complexation may be relatively transient as proposed recently (17).

As mentioned above, 5LO may become damaged or even fragmented after leaving the FLAP-disc. For the cases 5LO has not formed a complex, *e.g*. controls where 5LO and FND were mixed without additions or where complexes were never formed for other reasons, 5LO appears intact. On native gels as in the one in Figure 1 in the S2_file.pdf, lanes A1 and B2, a split of the 5LO was observed or a degradation product was present in the sucrose fractions (Fig. 3). On the native gels the split 5LO band Mw are still around 78 kDa whereas on the sucrose purified fractions, where the bands should only represent any of the three proteins, a breakdown product of the 5LOFNDCa complex is around 40 kDa. This implies an origin in the 5LO as intact FLAP and MSP are both smaller than 40 kDa.

*In vivo*, 5LO is a stable protein stored in the cytosol or in the nucleus depending on the cell type and rapidly shuttled between these compartments when needed. FLAP is found both in inner and outer nuclear envelope whereas LTC4 synthase is found in the latter only. It appears as if 5LO would have to be a relatively stable protein allowing storage until use and being shuttled. In both cytosol and nucleus, the environment is reducing, which is also necessary to maintain *in vitro* to prevent 5LO dimerization via cysteine bridges (11, 48). If degradation is induced in macrophages, the pathway involves ubiquitination (49). *In vitro*, inactivation of 5LO may be turn-over- or non-turnover-induced; the latter for which several measures must be taken to prevent this during purification of this Fe^2+^-containing protein in ambient oxygen (23). In particular, catalase and FeSO4 are present during purification to prevent the formation of oxygen radicals which could destroy the protein. However, these additions are later removed to avoid interference with the 5LO activity measurements. EDTA is kept in our buffers which reduces trace metal availability for radical formation due to Fenton reactions. In fact, FLAP has been described as a gatekeeper or helper and it was proposed that the N-terminal tail of 5LO carrying a FY-motif acts like a cork and regulator (15). In cells, non-bound 5LO is protected from non-turn-over inactivation by this cork and remains stable in the cells up to 4 hours with or without FLAP present in the membrane. However, upon recruitment to the membranes by Ca^2+^ addition, 5LO was rapidly inactivated by a turn-over dependent mechanism (15). It appeared that without or with Ca^2+^, 5LO is safe as long as FLAP is absent. With FLAP in the membranes, 5LO is affected to show lower concentration in the cytosol and really low presence if located on the membrane with FLAP due to turn-over based inactivation and destruction. In other words: For 5LO, presence of FLAP is problematic already in the absence of Ca^2+^ but worsens when located on the membrane/FLAP and turns over. It could be that upon 5LO binding to the FND, FLAP promotes the regulatory FY-cork to open up the active site for interaction with AA if it is present. In the case AA is not available since the complex was formed due to calcium binding, protein destruction could be initiated due to the oxygen present and the Fe^2+^ in 5LO being unprotected so that a Fenton reaction could take place producing hydroxyl radicals, *e.g*. Fig 3 left side. If AA is present in the FND-FLAP complex and presented to 5LO a more stable complex could initially form. The active site could be protected from radical formation for a longer period as a turn-over based inactivation may not occur *in vitro* as it appears that the intact nuclear membrane must be available (8) or it is due to the lack of the small Coactosin-like protein observed to bind on 5LO and increase activity (50).

In conclusion, the FND is a promising in vitro platform for studies of the 5LO-FLAP interaction. Future investigations of the cause for the observed 5LO damage could lead to insights into the 5LO-FLAP activation mechanism. Parameters to explore could be specific lipid compositions of the FND, presence of the Coactosin-like protein, test the combination of AA and Ca^2+^ and order of mixing all components as well as investigate the effect of inhibitors of either 5LO or FLAP. Ultimately changes in the leukotriene product profile related to specific parameters of the complex formation could lead to a mechanism for the very first step of AA conversion to leukotriene A4.

**S1 Raw images.**

**S2 File**: Figures 1 and 2. Results from representative examples of native gel electrophoresis in Figure 1 and 2D classifications of negatively stained (uranyl formate) samples in Figure 2.

## Acknowledgments

We acknowledge staff and resources at Protein Science Facility at Karolinska Institutet. In-gel digestion, peptide extraction, mass spectrometric analysis and database searches for protein identification were carried out at the Proteomics Facility, Biomedicum, Karolinska Institutet.

---

S1 raw files at the end.

**S2 File**: Figures 1 and 2 containing the results from representative examples of native gel electrophoresis and negative staining (uranyl formate) of the complexes.

**Figure 1.**
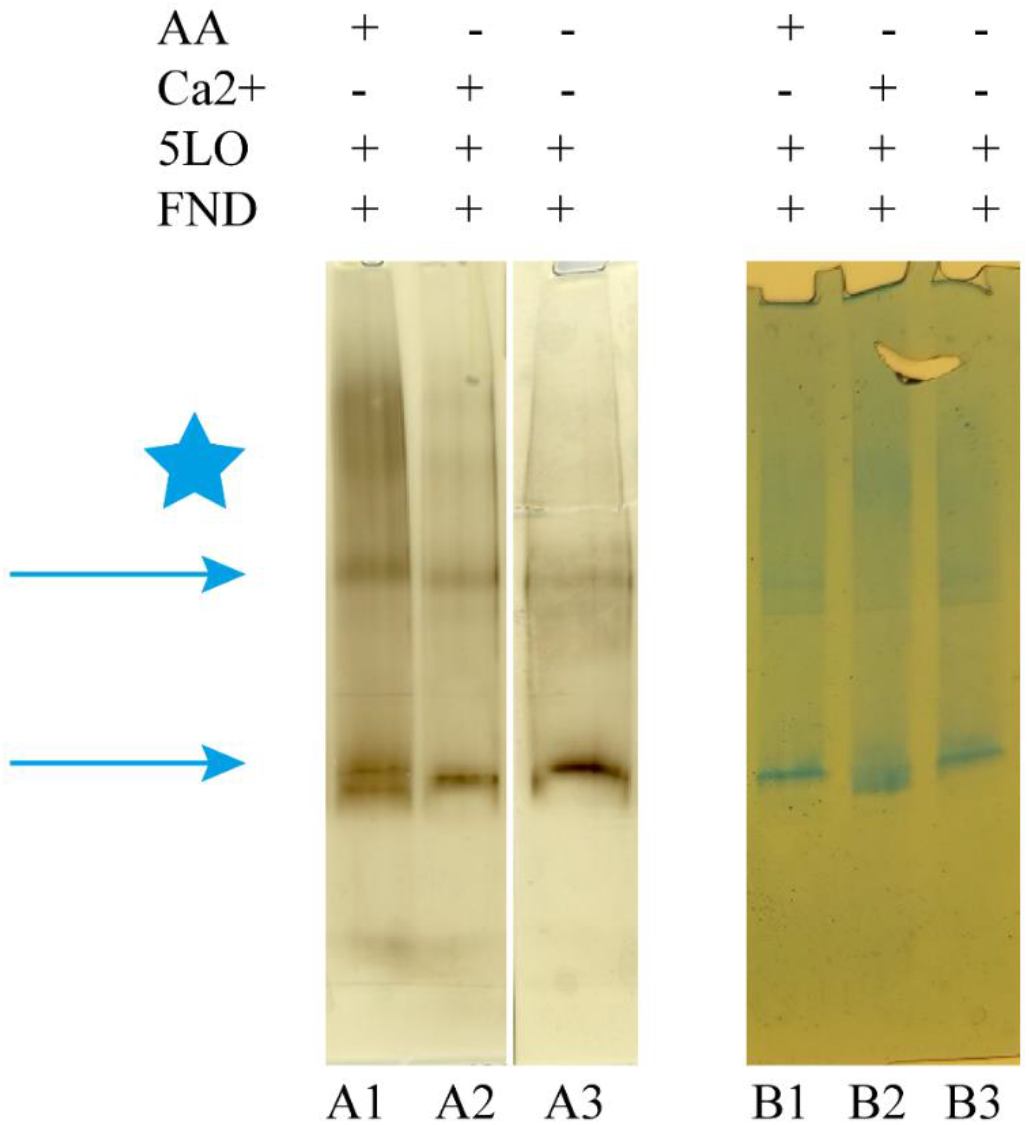
The presence of AA is sufficient to induce binding of 5LO on the FLAP-nanodiscs. **Formation of the complex 5LOFND, in the presence of either AA or Ca^2+^-ions.** The gels shown contained neither Ca^2+^ nor AA in the buffers during the gel-runs. Both gels were 4-16% Tris glycine Native page (A) or Blue Native page (B). The columns of + or – indicates the substances present in the sample mixture loaded in corresponding gel lane. The star indicates the level where bands show which may correspond to 5LOFND formed by the presence of either AA or Ca^2+^. To make the figure, relevant lanes were cut in the original gel images and rearranged in (A). A section was cut out from the original gel image in (B), see S1_Raw_files.pdf.

For the two gels in Figure 1, the sample preparations were different to test if the order of addition could have effects on 5LOFND complex-formation. For the gel in Fig. A, a specific amount of substance was added to the FND solution and incubated for 5 minutes before addition of 5LO in equimolar amounts to FND (0.6 μM) and left to react for another 5 minutes. The amount of substance added provided concentrations of 50 μM AA (lane A1) or 1 mM Ca^2+^ (lane A2) both in the presence of 0.5 mM EDTA.

This sample preparation is similar to that used for activity measurements earlier (Kumar *et al*., 2017). Under these conditions of substance additions the 5LOFNDAA complex formed (Lane A1) indicated by bands at higher levels (star) than FND, the FND band (upper arrow) was slowed down and a split 5LO band appears (lower arrow). The bands representing the Ca^2+^ induced complex appear weak (Lane A2, star-level) under these conditions.

A 5LOFNDCa complex was formed however, using another order of addition. For the gel in Fig. B, the samples were made by placing drops containing either AA, Ca^2+^ or just buffer on the wall of an Eppendorf tube followed by rapid centrifugation to mix the substance additions into the pre-mixed equimolar 5LO and FLAP solutions. The final concentrations are the same as above, 0.6 μM protein, 50 μM AA or 1 mM CaCl2. In Lane B2, the rapid addition of Ca^2+^ appears to have induced the formation of a 5LOFNDCa complex at a higher Mw (star) at the expense of the FND for which the band has almost disappeared (upper arrow). In addition, the 5LO band appears split (lower arrow, Lane B2). Lane B1, where the sample should contain the AA complex, displays a very weak band at higher Mw (Lane B1), the 5LO band is not split and the bands representing FND and 5LO have intensities similar to the control in lane B3.

**Figure 2.**
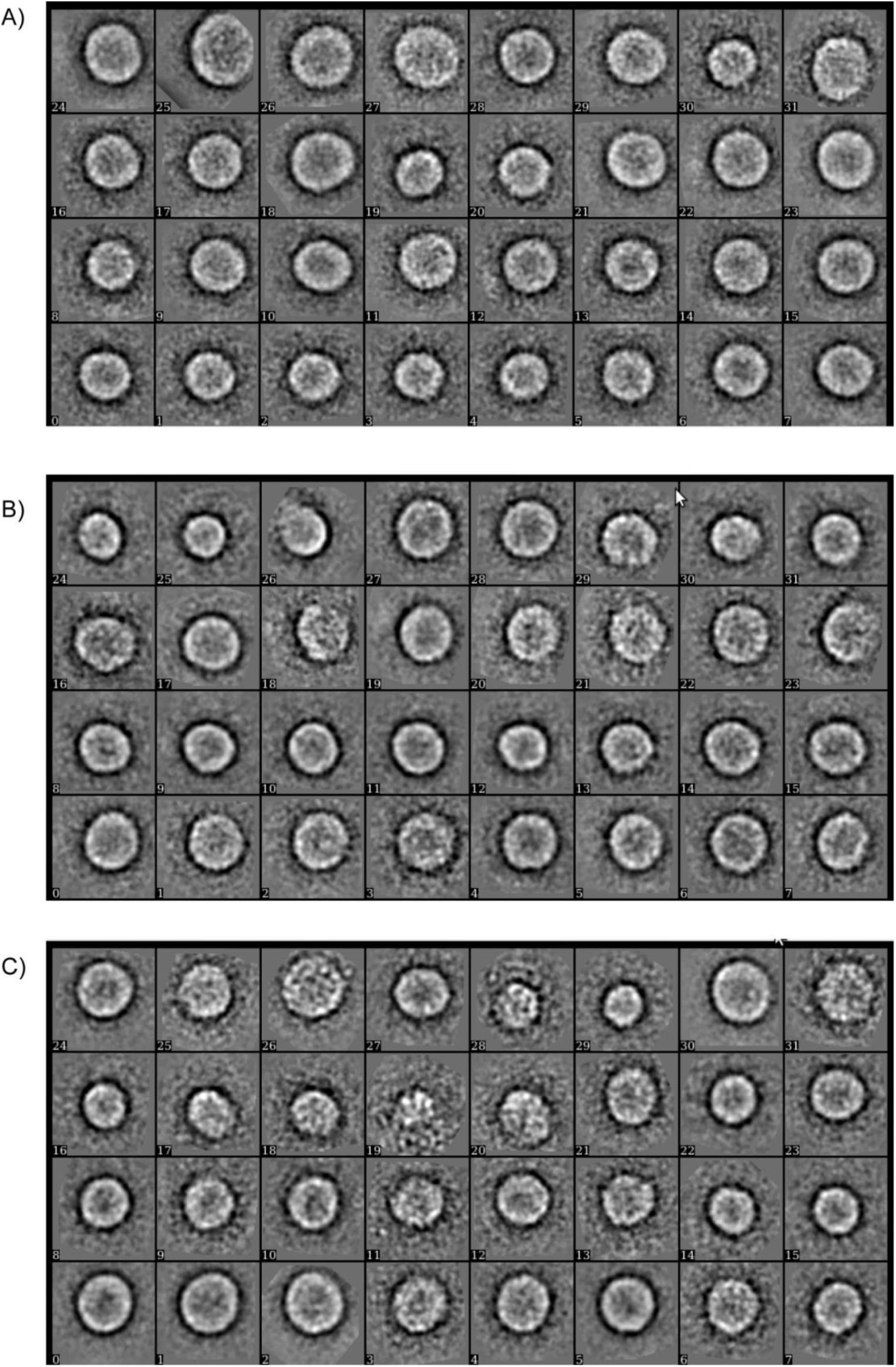
2D analysis of UF stained samples. 2D-class averages of data processed by EMAN2 (Tang *et. al*. 2007) for UF-stained samples: A) FND B) 5LOFNDAA and C) 5LOFNDCa. Particle sets were 3081 (FND), 3705 (5LOFNDAA) and 3201 (5LOFNDCa). The box side length is 30.1 nm.

## S1_Raw_images

**Main text Fig 1A:**
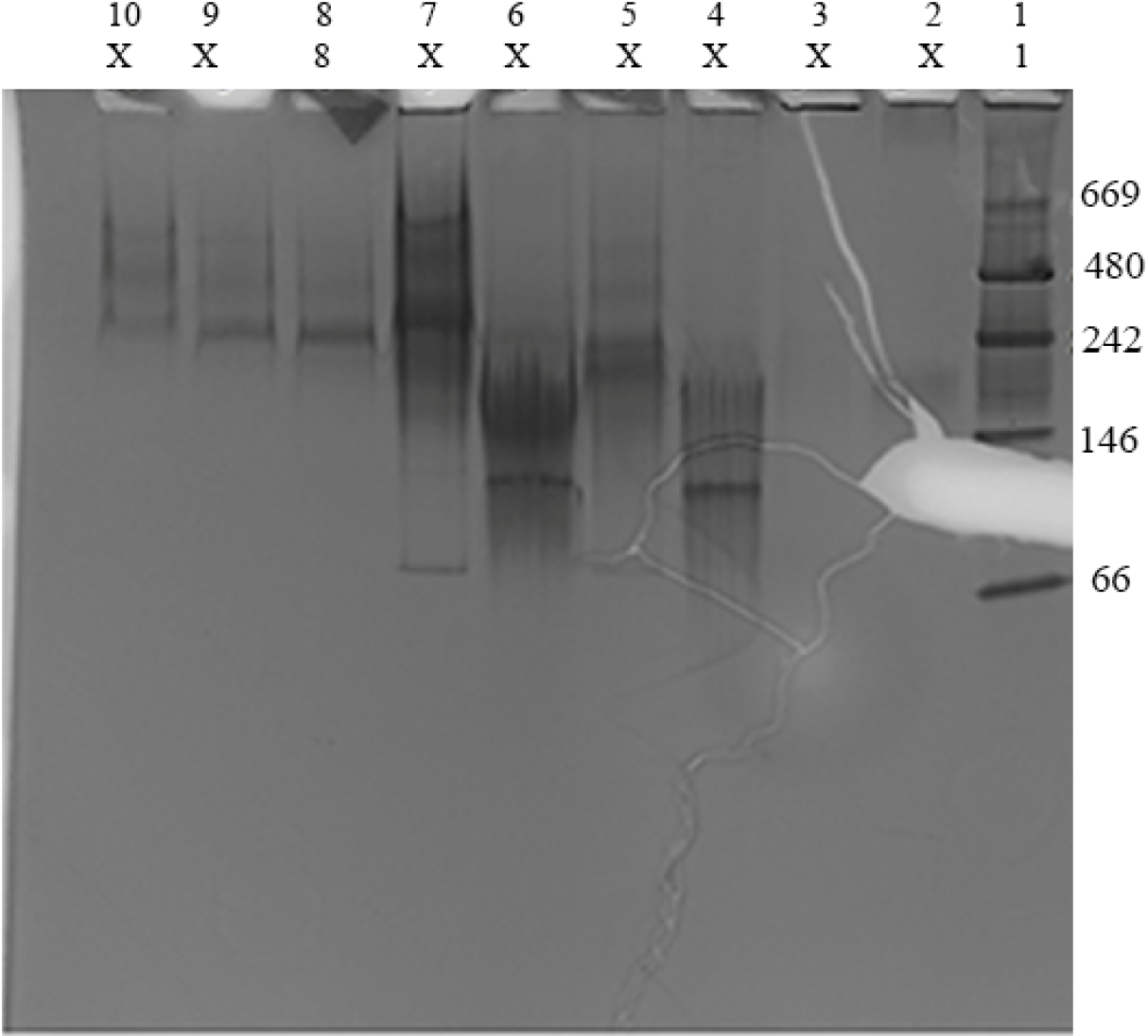
The marker in Lane 1 is to the right in both gels. Purified nanodiscs containing FLAP shows in Lane 8 in the original gel and to the left of the marker in Main text Fig 1A. The original gel shows most steps in the reconstitution procedure of FLAP into nanodiscs. Lanes 8-10 are elution fractions from a size-exclusion column purification. The gel is a 4−20 % tris-glycine native PAGE.

**Main text Fig 1B:**
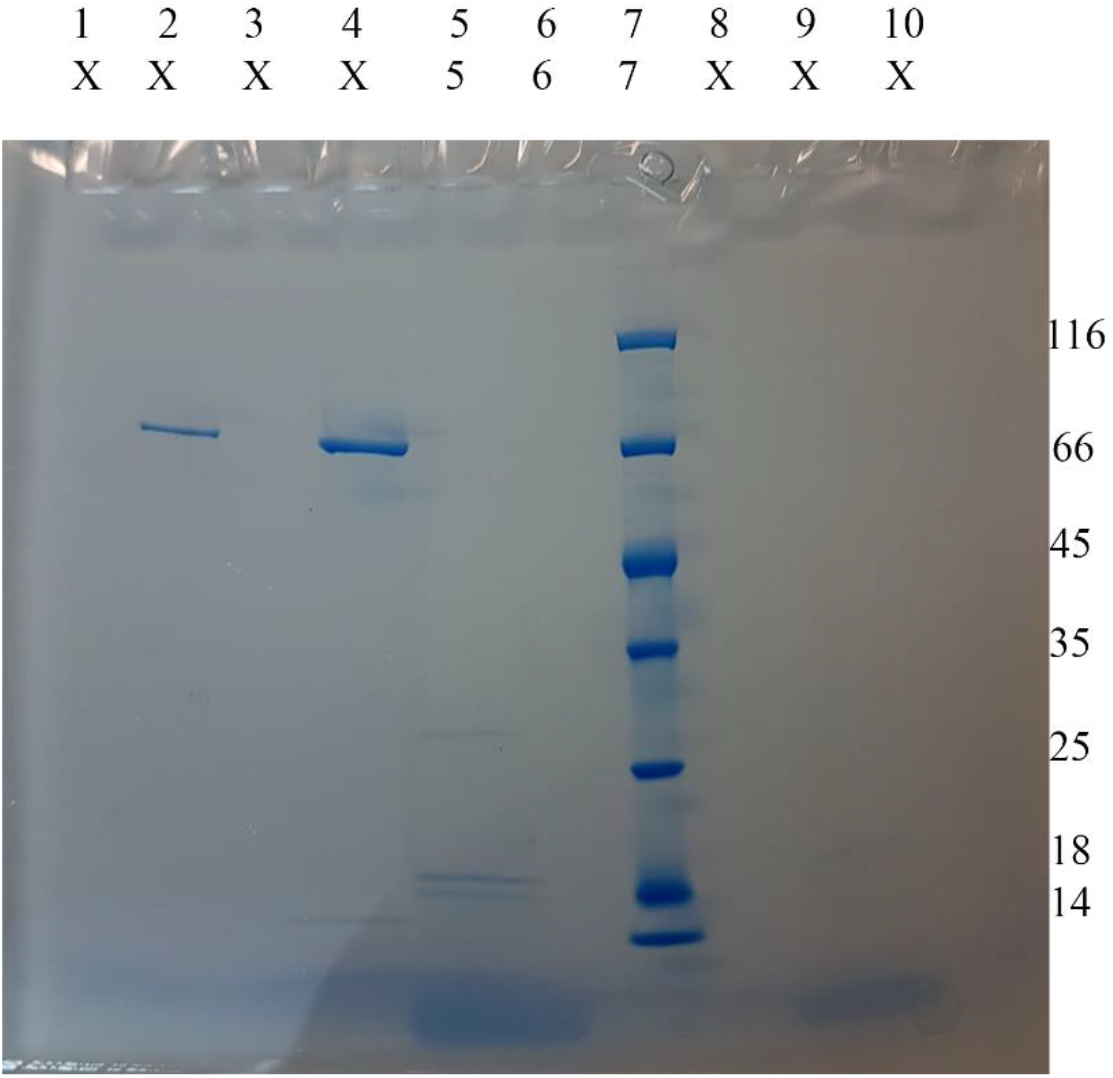
The marker in Lane 7 here is to the right in Main Fig 1B. Lane 6 is empty. Lane 5 comprises nanodiscs containing FLAP. Denaturing by SDS results in bands from MSP 1E3D1, ca 25 kDa, and FLAP-C-His6, 18.5 kDa. The band at ca 18 kDa could indicate a breakdown product of FLAP. Cropped parts of Lanes 5-7 are present in Main text Fig 1B. Lanes 2 and 4 shows 5LO at different concentrations. Lanes 1, 3, 6, 8 and 10 are empty. The gel is a 10-20 % tris-glycine SDS PAGE

**Supplementary Fig 1A:**
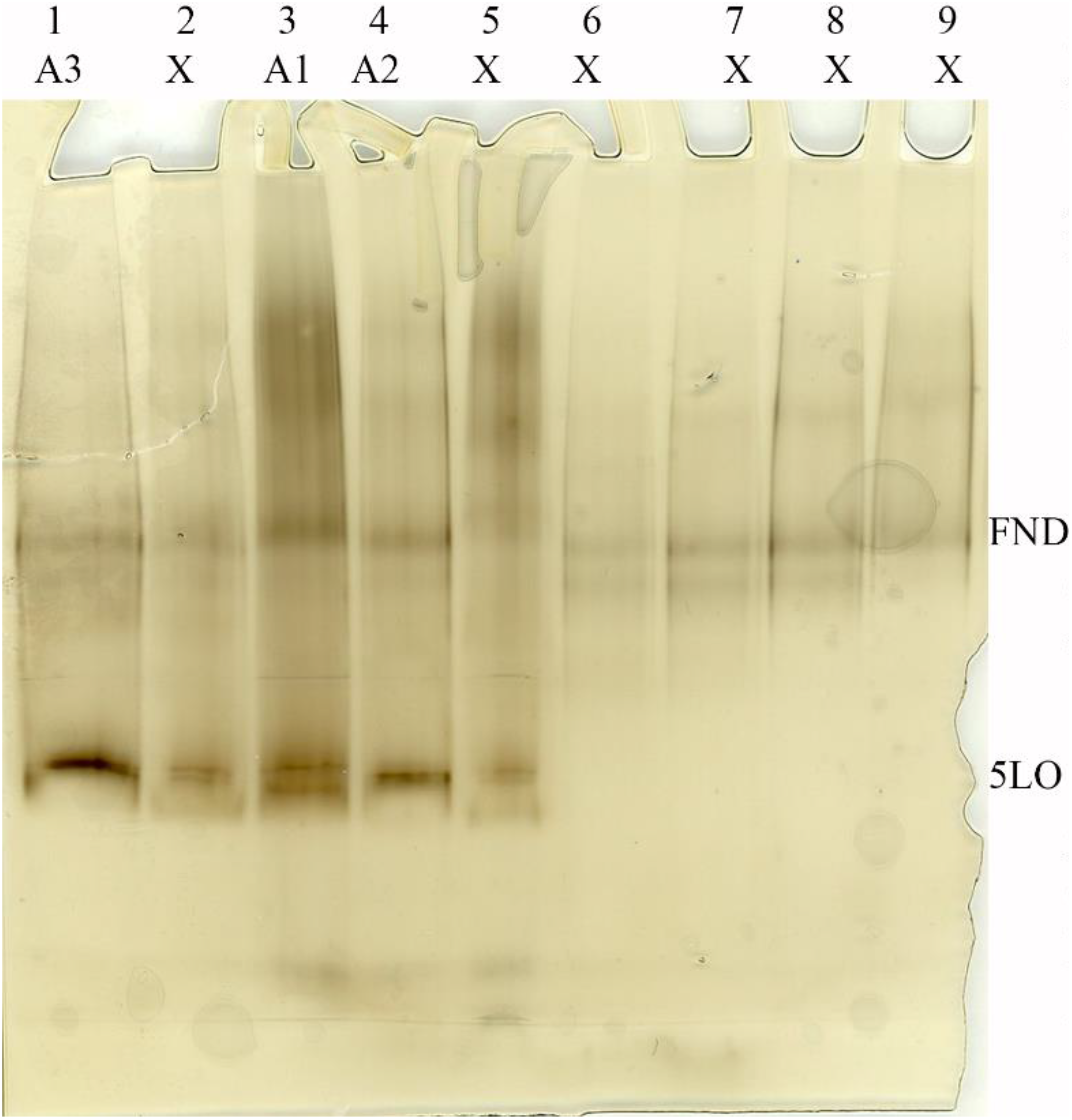
The gel was run without marker. FND (ca 242 kDa) and 5LO (78 kDa) served as markers. Lanes 1-3 were rearranged to lanes A1-A3 in S1 file Fig 1A for comparison to S1 file Fig 1B. The samples loaded were:

A1 the AA5LOFND complex
A2 the Ca5LOFND complex
A3 the 5LO and FND w/o adds The gel is a 4-16 % tris-glycine native PAGE, silver stained. No AA or Calcium were present during the gel-run. Therefore part of the complexes formed before loading separates into free 5LO and FND

**Supplementary Fig 1B:**
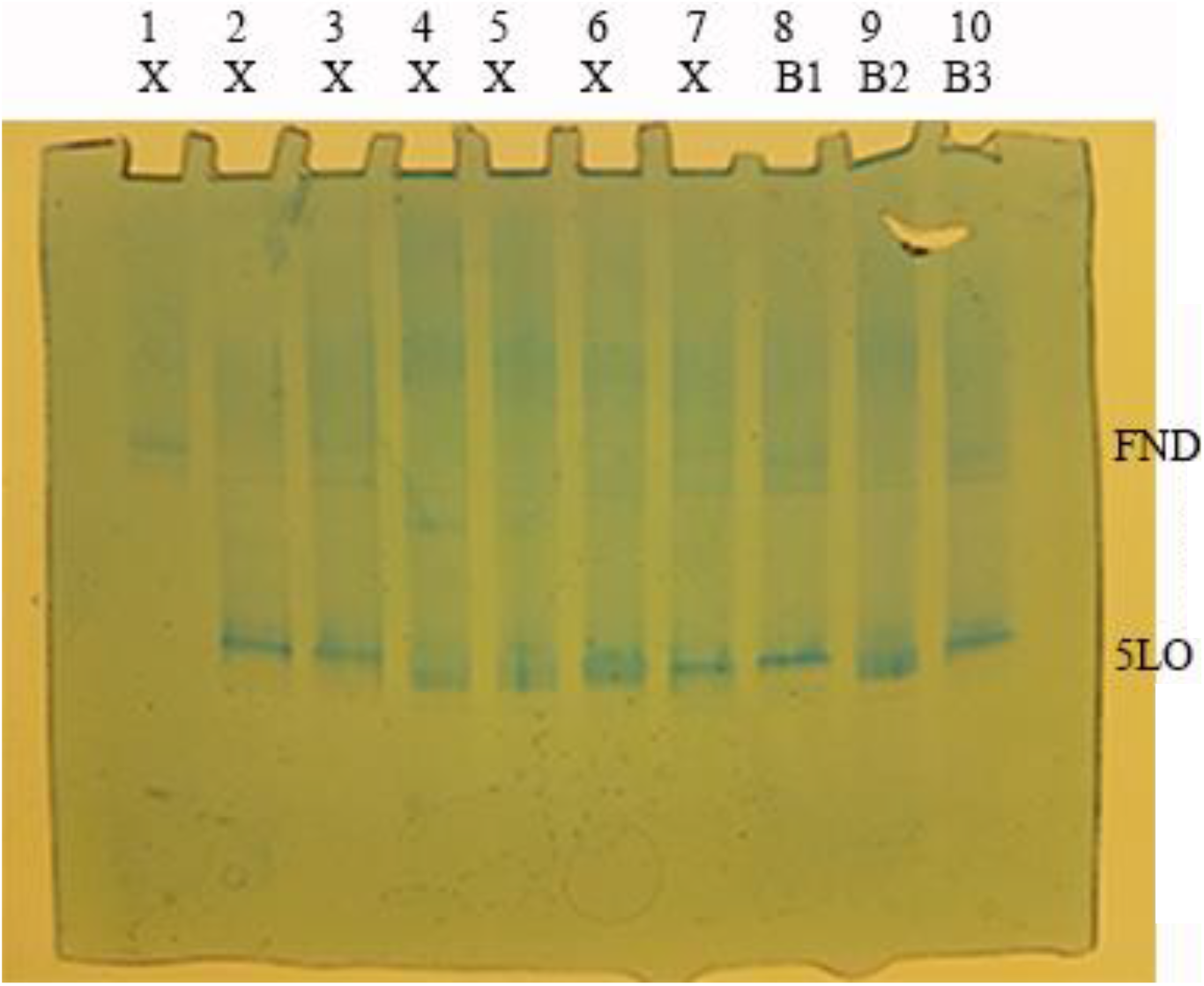
The gel was run without marker. FND (ca 242 kDa) and 5LO (78 kDa) served as markers. Lanes 8-10 were renamed to lanes B1-B3 in S1 file Fig1B. The samples loaded were:

B1 the AA5LOFND complex
B2 the Ca5LOFND complex
B3 the 5LO and FND w/o adds The gel is a 4-16 % tris-glycine Blue Native PAGE. An arte-factual “band” is located below the FND. No AA or Calcium were present during the gel-run. Therefore part of the complexes formed before loading separates into free 5LO and FND.

**Main text Fig 3:**
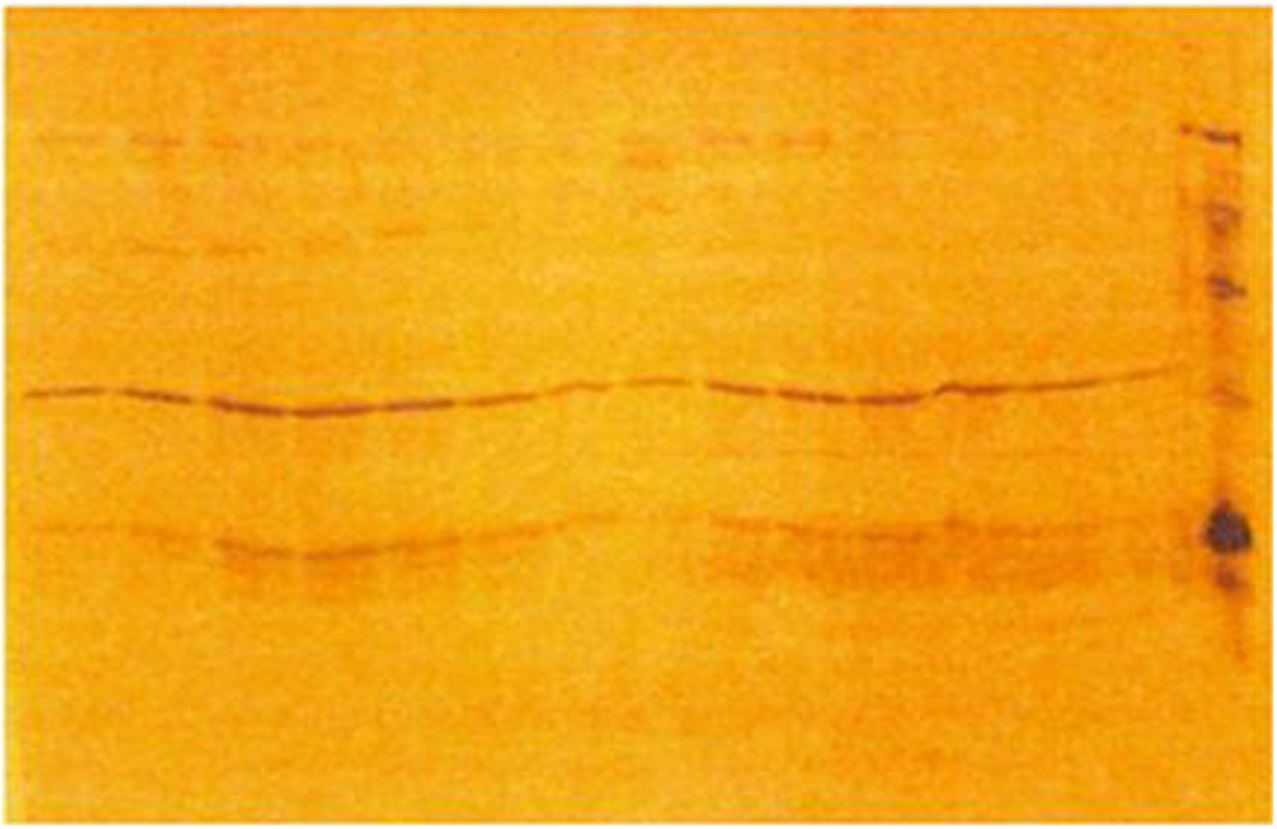
The original gel is presented the main text in Fig3.

